# An in vivo rat lumbar spine instability model induced by intervertebral disc injury

**DOI:** 10.1101/2025.05.14.653956

**Authors:** Fangxin Xiao, Wendy Noort, Juliette Lévénez, Jia Han, Jaap H. van Dieën, Huub Maas

## Abstract

Intervertebral disc (IVD) degeneration is a potential contributor to low-back pain. While experimental IVD injury models have demonstrated IVD structural changes, the early mechanical consequences remain unclear. This study aimed to establish a rat model of lumbar spine instability via IVD injury and assess back musculature adaptations to IVD injury. Thirty-one adult male Wistar rats were assigned to three groups: IVD knife stab lesion (knife), IVD needle puncture (needle), and sham surgery control (control). In the knife and needle groups, L4/L5 IVDs were injured at 14 weeks of age. One-two weeks post-intervention, lumbar multifidus (MF) and medial longissimus (ML) muscles were excised, L4-L5 spinal segments were harvested for mechanical testing, and IVDs were collected for histology. The needle group exhibited lower peak stiffness, peak moment, and hysteresis than controls in flexion, with no difference in lateral bending. IVD height and area did not differ between groups, but the needle group had a smaller nucleus relative to annulus area compared to controls. Morphological changes were observed in both injury groups. The needle group showed higher normalized ML mass, while normalized MF mass was unchanged. In conclusion, a rat model of lumbar spine instability was successfully established via IVD needle injury.

## Introduction

Low-back pain (LBP) is a leading cause of years lived with disability worldwide (Hartvigsen et al. 2018) and spinal instability may be one of the contributing mechanisms (Panjabi 2003). Spinal instability is clinically defined as hypermobility under physiological loads (Farfan and Gracovetsky 1984), which has been observed in people with LBP (Fujiwara, Lim, et al. 2000; Passias et al. 2011) and may lead to altered motor control and load sharing among spinal structures (van Dieen, Selen, Cholewicki 2003; van Dieen et al. 2019).

IVD degeneration is significantly correlated to instability (Knutsson 1944; Fujiwara, Lim, et al. 2000; Fujiwara, Tamai, et al. 2000; Tanaka et al. 2001; Kong et al. 2009; Galbusera et al. 2014; Swanson and Creighton 2020), shown as increased segmental mobility with increasing severity of IVD degeneration up to Grade IV (Fujiwara, Lim, et al. 2000; Tanaka et al. 2001). Experimental IVD disruption has been used to induce IVD degeneration in various animal models (Jin, Balian, Li 2018; Desmoulin, Pradhan, Milner 2020), especially in rodents (Shi et al. 2018). Mechanical injury models, including depressurization of the nucleus pulposus (NP) and/or disruption of the annulus fibrosus (AF), for which scalpel stabbing or needle puncturing are most frequently used (Alini et al. 2008; Jin, Balian, Li 2018), allow precise control over the severity and timing of the injury (Alini et al. 2008).

Physical disruption of the IVD structure has been demonstrated to induce rapid IVD degeneration. Inflammatory responses (Ulrich et al. 2007), increased catabolic (Chen CH et al. 2016) and fibrotic activity (Glaeser et al. 2020), and downregulated main NP matrix gene and aggrecan gene expression levels (Glaeser et al. 2020) were detectable one week after injury. MRI imaging of rat tail IVD confirmed severe microstructural degeneration, reflected in diffusion properties, one week after IVD puncture (Li et al. 2019). Furthermore, morphological changes (Rousseau et al. 2007; Ulrich et al. 2007; Korecki, Costi, Iatridis 2008; Chen T et al. 2018; Maas et al. 2018; Mosley et al. 2020), reduced NP chondrocytes (Chen T et al. 2018), and increased apoptosis (Korecki, Costi, Iatridis 2008) were found. Injured discs exhibited decreased height (Kim et al. 2011; Lai et al. 2016), a disorganized AF, decreased NP size (Rousseau et al. 2007; Maas et al. 2018), and blurring of the interface between AF and NP. These molecular and morphological changes may affect the IVD’s mechanical properties (Yerramalli et al. 2007; Iatridis et al. 2013; Fontana, See, Pandit 2015).

However, our understanding of the mechanical consequences of IVD injury is limited. In rat, in vitro studies have shown that IVD injury acutely compromised bending (Xiao et al. 2023) and torsional (Wang et al. 2022) mechanical properties. In vivo studies showed that, despite the presence of structural degenerative changes, IVD mechanics were not affected by injury, neither after four (Rousseau et al. 2007), nor after six weeks (Mosley et al. 2020).

The mechanical stability of the spine is critical to its proper functioning, for example, to deal with instantaneous changes in spinal posture (Panjabi 1992). When there is dysfunction in one of the spinal stabilizing subsystems, adaptations in one or more of the other spinal stabilizing subsystems (e.g., musculotendinous subsystem) may occur (Panjabi 1992). For instance, decreased multifidus muscle (MF) mass has been observed at one week after rat lumbar IVD injury, while longissimus muscle (ML) mass tended to increase (Maas et al. 2018). Thus, an understanding of the short-term effects of IVD injury on its mechanical properties, as well as the link between structural and mechanical changes, is required to better evaluate the (neuro)mechanical effects of IVD injury.

This study aimed to (1) establish a rat model of lumbar spine instability by damaging the lumbar IVD, (2) assess adaptations of back musculature to IVD injury. We hypothesized that (1) IVD injury changes bending mechanics of the spinal segments, including reduced peak stiffness, peak moment, and hysteresis. (2) IVD injury will cause atrophy of MF, while ML mass will increase or remain unchanged.

## Materials and Methods

### Animals

A total of 31 adult male Wistar rats (*Rattus norvegicus*) were assigned to three groups: IVD knife stab lesion (knife, n=8), IVD needle puncture (needle, n=14), and sham surgery control (control, n=9). The rats in the knife and control groups were randomly assigned within one study, with sample size determined by a prior power analysis (power=0.8, alpha=0.5) using G*Power (Faul et al. 2007), while those in the needle group came from a separate study. Surgical and experimental procedures were in agreement with the guidelines and regulations concerning animal welfare and experimentation set forth by Dutch law on animal research in full agreement with the Directive 2010/63/EU, with local approval by and under supervision of the local Animal Welfare Body, and approved by the Central Commission for Animal Experiments of the Netherlands Government at the Vrije Universiteit Amsterdam (Permit Number: AVD11200202115388).

### Surgical Procedures for IVD injury

Body mass at surgery and sacrifice are reported in Table 1. IVDs were injured when the rats were 14 weeks old. Carprofen (3 mg/kg, Rimadyl®, Zoetis B.V., Capelle a/d Ijssel, The Netherlands) was administrated subcutaneously 12 h before surgery. Carprofen and buprenorphine (0.02 mg/kg, Buprecare®, Ecuphar NV, Oostkamp, Belgium) were administrated subcutaneously 30-60 min before surgery. The rats were anaesthetized using isoflurane (induction: 3-5%, maintenance: 1-2%) and placed on a heating pad. A few minutes before skin incision, ropivacaine (2 mg/kg, Fresenius Kabi Norge AS, Halden, Norway) was injected in the skin. All surgeries were performed under aseptic conditions. Body temperature, breathing rate and heart rate, anaesthetic depth were monitored every 10-30 min. Two days post-surgery, carprofen was administrated subcutaneously. Three-five days post-surgery, carprofen was provided in drinking water (0.06 mg/ml) and mixed into food. After surgery, the rats were allowed to move freely in their cage with access to food and water *ad libitum* at a 12-hour day-night cycle. Body mass changes were monitored for five days. The rats were housed in pairs, except for the first 24 hours post-surgery, when the two rats in the same cage were separated by a cage divider.

**Table 1.** Body mass, muscle mass and IVD histology.

|  |  | Body<br>Mass<br>(T0)<br>(gram) | Body<br>Mass<br>(T1)<sup>&#</sup><br>(gram) | IVD<br>Height<br>(mm) | IVD<br>area<br>(mm<sup>2</sup>) | NP<br>area<br>(mm<sup>2</sup>) | Relative<br>NP<br>area<sup>#</sup> | Normalized<br>MF mass | Normalized<br>ML mass |
| --- | --- | --- | --- | --- | --- | --- | --- | --- | --- |
| sample<br>size | control | 9 | 9 | 9 | 9 | 9 | 9 | 8 | 7 |
|  | knife | 8 | 8 | 8 | 8 | 8 | 8 | 8 | 8 |
|  | needle | 14 | 14 | 13 | 9 | 9 | 9 | 14 | 14 |
| mean<br>(SD) /<br>median<br>(IQR) | control | 406<br>(56) | 442<br>(51) | 1.38<br>(0.23) | 9.43<br>(2.02) | 1.09<br>(0.278) | 0.128<br>(0.073) | 0.00116<br>(0.00011) | 0.00185<br>(0.00011) |
|  | knife | 419<br>(73) | 440<br>(47) | 1.27<br>(0.26) | 10.3<br>(1.87) | 1.01<br>(0.388) | 0.089<br>(0.057) | 0.00113<br>(0.00015) | 0.00188<br>(0.00038) |
|  | needle | 356<br>(34) | 373<br>(26) | 1.28<br>(0.26) | 10.2<br>(1.48) | 0.562<br>(0.46) | 0.048<br>(0.089) | 0.00108<br>(0.00012) | 0.00222<br>(0.00030) |

*T0, Body Mass at time of surgery; T1, Body Mass at time of sacrifice; <sup>&</sup>Body Mass at time of sacrifice was used for correlation analysis; <sup>#</sup>Non-normally distributed data. Data are presented as mean (SD) for normally distributed data and median (IQR) for non-normally distributed data. IVD, intervertebral disc; NP, nucleus pulposus; MF, multifidus muscle; ML, longissimus muscle*

In injury groups, the L4/L5 IVD was injured using a transperitoneal-ventral approach as described previously (Fig.1A) (Maas et al. 2018). The IVD was injured ventral-dorsally in the middle to a depth of 2.5 mm fully penetrating the NP, using a scalpel blade (Swann Morton, SM61, max. width 1.6 mm, max. thickness 0.6 mm) or a 21-gauge hypodermic needle (Becton Dickinson, outer diameter 0.8mm) with a 360° rotation. Both the scalpel and needle had stoppers set at a depth of 2.5mm to ensure consistent penetrating depth. Subsequently, the abdominal wound and skin were closed with absorbable sutures (5-0, Vicryl, absorbable, ETHICON). Tiny dots of tissue glue (Vetbond™ Tissue Adhesive, 3M, Neuss, Germany) were applied between the skin sutures. The actual spine level of the injured IVD was confirmed at time of tissue extraction. In the control group, sham surgery included all procedures necessary to access the L4/L5 IVD, but no IVD injury was induced.

**Fig. 1.**
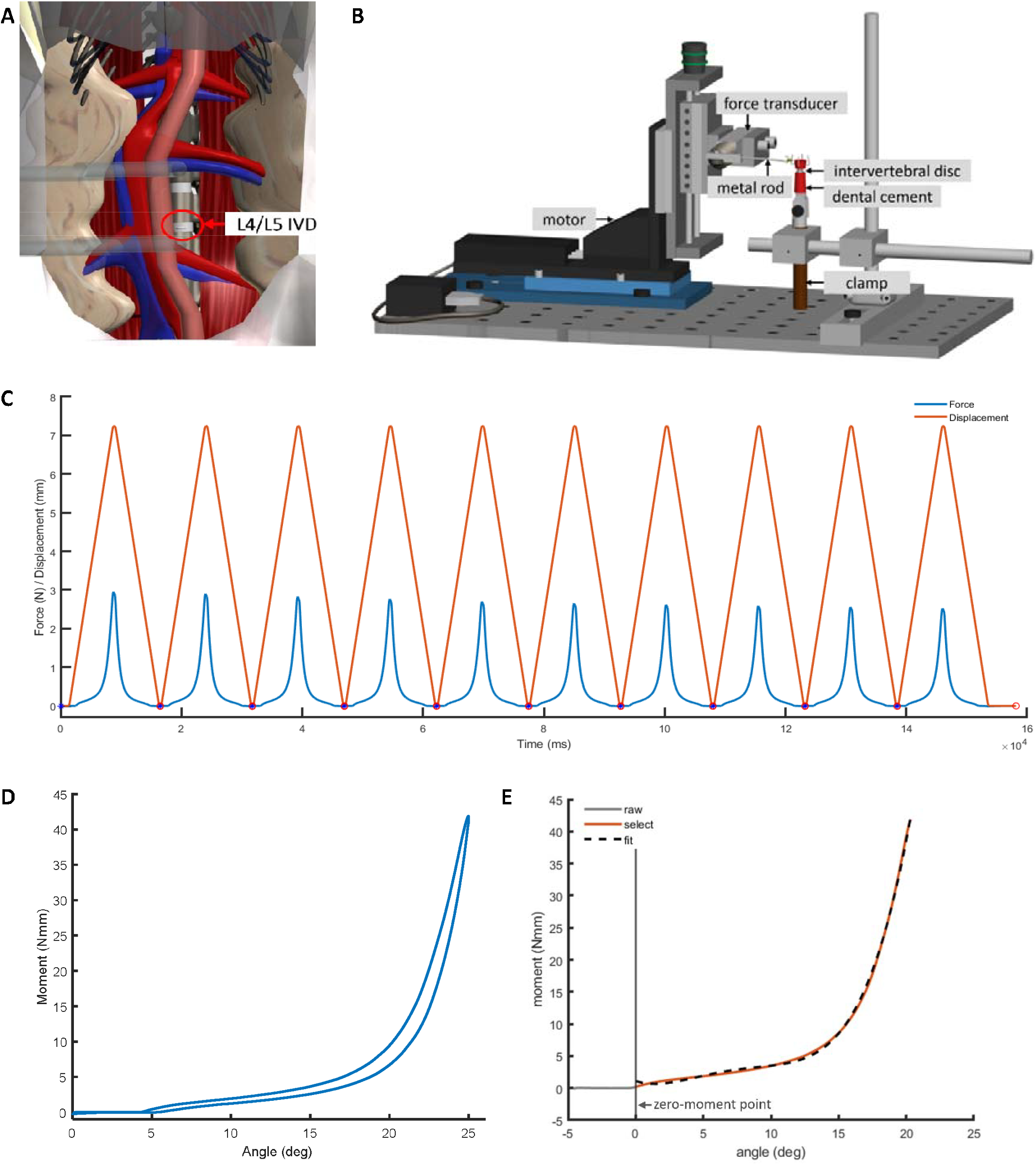
**(A)** Schematic of intervertebral disc injury surgical approach. Drawing was made by Guus Baan. **(B)** Mechanical test setup. Drawing was made by Guus Baan. **(C)** An example of filtered time series of force (blue) and displacement (red) signals from a flexion test. **(D)** Angle-moment curve of the 10th cycle from above signals. **(E)** loading part of the 10th angle-moment curve with adjusted ‘zero-displacement point’. Grey, raw data; red, selected data after removing the slack angle; black dashed line, 4th order polynomial fitted data.

### Tissue extraction

Rats in the knife and needle groups were sacrificed one week after surgery, those in the control group were sacrificed two weeks after sham surgery. 30-60 min prior to extraction of tissues, buprenorphine was administrated subcutaneously. Then the rats were anaesthetized using isoflurane and sacrificed with an overdose of pentobarbital (intracardial, Euthasol 20%, AST farma, Oudewater, The Netherlands). Immediately before (under isoflurane anaesthesia) or within two hours after sacrifice, left and right multifidus (MF) fascicles originating from L3-L4 vertebrae, as well as the medial longissimus muscles (ML) between L1-S3 were excised. The mass of the extracted muscles was measured, averaged across sides, and normalized to body mass. L4-L5 spinal segments (vertebra-IVD-vertebra) were collected and wrapped by saline-soaked gauze after removing soft tissues (muscles, supraspinous and interspinous ligaments), then stored at -20°C.

### Mechanical Testing

As described previously (Xiao et al. 2023), specimens were thawed at room temperature one day before mechanical testing and potted in dental cement (DuraLay, Reliance Dental Manufacturing, LLC, United States), then wrapped by phosphate-buffered saline (PBS)-soaked gauze and stored at +4°C in sealed double plastic bags. On the day of testing, the specimens were placed at room temperature in sealed bags for 1-4 h before testing.

Non-destructive displacement-controlled bending tests of the specimens were performed in a fixed sequence: flexion, left bending, and right bending (Fig.1B). Maximum bending angles were 25° for flexion and 15° for lateral bending. The motor moved at a constant velocity of 1mm/s, with a maximum acceleration/deceleration of 4.5 mm/s^2^. Custom LabVIEW software (LabVIEW 13.0, National Instruments, Austin, TX) controlled motor movement and collected displacement and force signals. For each direction, 10 cycles of testing followed a 10mN preload and 10 cycles of preconditioning. In between tests, the IVD was kept moist by wrapping the IVD with PBS-soaked gauze and spraying it with PBS every 2-3 min.

### Histology

Following mechanical testing, the IVDs were isolated from the spinal segments, frozen in liquid nitrogen and stored at -80°C. IVDs were sectioned transversally at 12 µm thickness in a cryostat at -20°C (Maas et al. 2018). Slides from the middle portion of the IVD were stained with Picrosirius Red (PR). From scanned images (20x magnification), the areas of the IVD and the NP were measured in ImageJ (National Institutes of Health, United States) by an assessor blinded to intervention. The relative NP area was calculated as the ratio of NP to IVD area.

### Data Analysis

Force and displacement signals were sampled at 100 Hz and second order, 5Hz, low pass zero-lag Butterworth filtered (Fig.1C). Angle-moment data of the 10^th^ cycle were used for data analysis (Fig.1D), hysteresis was calculated as the area between loading and unloading curves. Subsequently, the zero-moment point was identified by removing the slack angle after preconditioning and a fourth order polynomial was fitted to the loading part of the angle-moment curve (Fig.1E) (Xiao et al. 2023). Per direction, the lowest maximum angle across all specimens (19° for flexion, 9° for left and right bending after adjustment for zero-moment point) was used to calculate peak moment from the fitted angle-moment curve and peak stiffness from the analytical derivative of the fitted curve. All mechanical outcomes were normalized to body mass, as this was correlated to most of these outcomes (Table 2). Parameter calculation was conducted in random order by an assessor blinded to intervention.

**Table 2.**
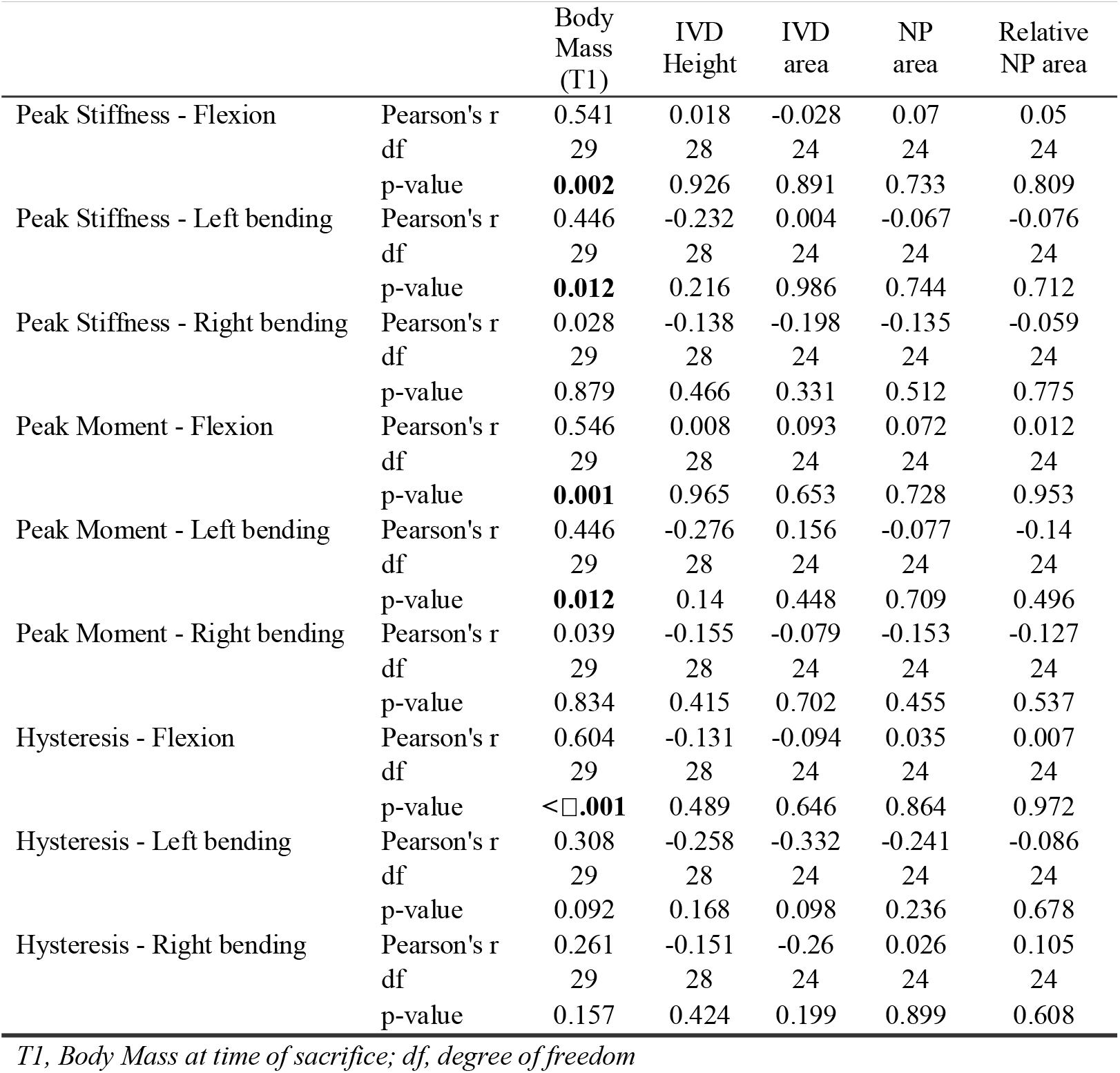
Correlation between Body Mass, IVD histology and IVD mechanical properties.

|  |  | Body<br>Mass<br>(T1) | IVD<br>Height | IVD<br>area | NP<br>area | Relative<br>NP area |
| --- | --- | --- | --- | --- | --- | --- |
| Peak Stiffness - Flexion | Pearson's r | 0.541 | 0.018 | -0.028 | 0.07 | 0.05 |
|  | df | 29 | 28 | 24 | 24 | 24 |
|  | p-value | **0.002** | 0.926 | 0.891 | 0.733 | 0.809 |
| Peak Stiffness - Left bending | Pearson's r | 0.446 | -0.232 | 0.004 | -0.067 | -0.076 |
|  | df | 29 | 28 | 24 | 24 | 24 |
|  | p-value | **0.012** | 0.216 | 0.986 | 0.744 | 0.712 |
| Peak Stiffness - Right bending | Pearson's r | 0.028 | -0.138 | -0.198 | -0.135 | -0.059 |
|  | df | 29 | 28 | 24 | 24 | 24 |
|  | p-value | 0.879 | 0.466 | 0.331 | 0.512 | 0.775 |
| Peak Moment - Flexion | Pearson's r | 0.546 | 0.008 | 0.093 | 0.072 | 0.012 |
|  | df | 29 | 28 | 24 | 24 | 24 |
|  | p-value | **0.001** | 0.965 | 0.653 | 0.728 | 0.953 |
| Peak Moment - Left bending | Pearson's r | 0.446 | -0.276 | 0.156 | -0.077 | -0.14 |
|  | df | 29 | 28 | 24 | 24 | 24 |
|  | p-value | **0.012** | 0.14 | 0.448 | 0.709 | 0.496 |
| Peak Moment - Right bending | Pearson's r | 0.039 | -0.155 | -0.079 | -0.153 | -0.127 |
|  | df | 29 | 28 | 24 | 24 | 24 |
|  | p-value | 0.834 | 0.415 | 0.702 | 0.455 | 0.537 |
| Hysteresis - Flexion | Pearson's r | 0.604 | -0.131 | -0.094 | 0.035 | 0.007 |
|  | df | 29 | 28 | 24 | 24 | 24 |
|  | p-value | **<0.001** | 0.489 | 0.646 | 0.864 | 0.972 |
| Hysteresis - Left bending | Pearson's r | 0.308 | -0.258 | -0.332 | -0.241 | -0.086 |
|  | df | 29 | 28 | 24 | 24 | 24 |
|  | p-value | 0.092 | 0.168 | 0.098 | 0.236 | 0.678 |
| Hysteresis - Right bending | Pearson's r | 0.261 | -0.151 | -0.26 | 0.026 | 0.105 |
|  | df | 29 | 28 | 24 | 24 | 24 |
|  | p-value | 0.157 | 0.424 | 0.199 | 0.899 | 0.608 |

*T1, Body Mass at time of sacrifice; df, degree of freedom*

### Statistical Analysis

MATLAB R2021a (MathWorks, Inc., Natick, MA, United States) was used for statistical analysis. Normality was tested with the Shapiro-Wilk test; homogeneity of variance was tested with Levene’s test (Appendix A). Pearson correlation coefficient was calculated to determine associations between body mass, IVD height, IVD area, NP area, relative NP area, and IVD mechanics. The fits to loading parts of the angle-moment curves of the 10^th^ testing cycle of all IVDs were compared between groups using the MATLAB based spm1d-package (SPM) (Pataky 2010). According to data distribution, parametric or non-parametric tests *“ex1d_anova1*” were used. Differences were considered significant if any SPM values exceeded the critical threshold. Depending on data distribution, parametric (one-way ANOVA with Tukey adjusted pairwise comparisons) or non-parametric (Kruskal-Wallis with Mann-Whitney U test pairwise comparisons) tests were used, to evaluate differences between groups for IVD structural and mechanical properties, and muscle mass. For all comparisons, alpha was set at 0.05. Data are presented as mean (SD) or median (IQR).

## Results

During the recovery period, several rats had chewed the skin incisions and sutures open, local wound were treated and re-sutured immediately after observation. During tissue extraction, we found in the needle group, injuries at L3/L4 in two and at L5/L6 in four rats. In those cases, the corresponding spinal segments were collected for mechanical and histological analysis. Body mass at termination was significantly lower in the needle group than in the control and knife groups (*p*=0.015, χ^2^_(2)_=8.384) (Table 1).

### IVD injury induced IVD mechanical changes

No significant group differences were found in angle-moment curves in flexion, but significant differences were found in left bending in the initial 6% of the bending angle (*p*=0.040, F=6.133) (Appendix B, Fig.1.B.1-2) and in right bending in the initial 1.4% (*p*=0.040, F=5.507) (Appendix B, Fig.1.C.1-2), where the needle group showed higher moments than the control group and/or the knife group. However, moments in this toe-region of the angle-moment curves were very low, approaching zero. Hence, we do not consider this to be functionally relevant.

IVD mechanical properties in flexion differed significantly between groups (Fig.2 & Table 3-5). (1) Slack angle was smallest in the knife group, showing a 36% (1.5°) reduction compared to the control group (*p*=0.022, t=2.850). (2) Peak stiffness differed between groups (*p*=0.011, F_(2,28)_=5.348), with the needle group having a 54% lower stiffness than the control group (*p*=0.008, t=3.260). (3) Peak moment varied significantly among groups (*p*=0.024, χ^2^_(2)_ =8.657), with the needle group showing a 55% lower moment than the control group (*p*=0.006, U=107). (4) Hysteresis also differed between groups (*p*=0.036, F_(2,28)_=3.735), but pairwise comparisons showed no significant differences. No significant differences were found between groups in left or right bending.

**Fig. 2.**
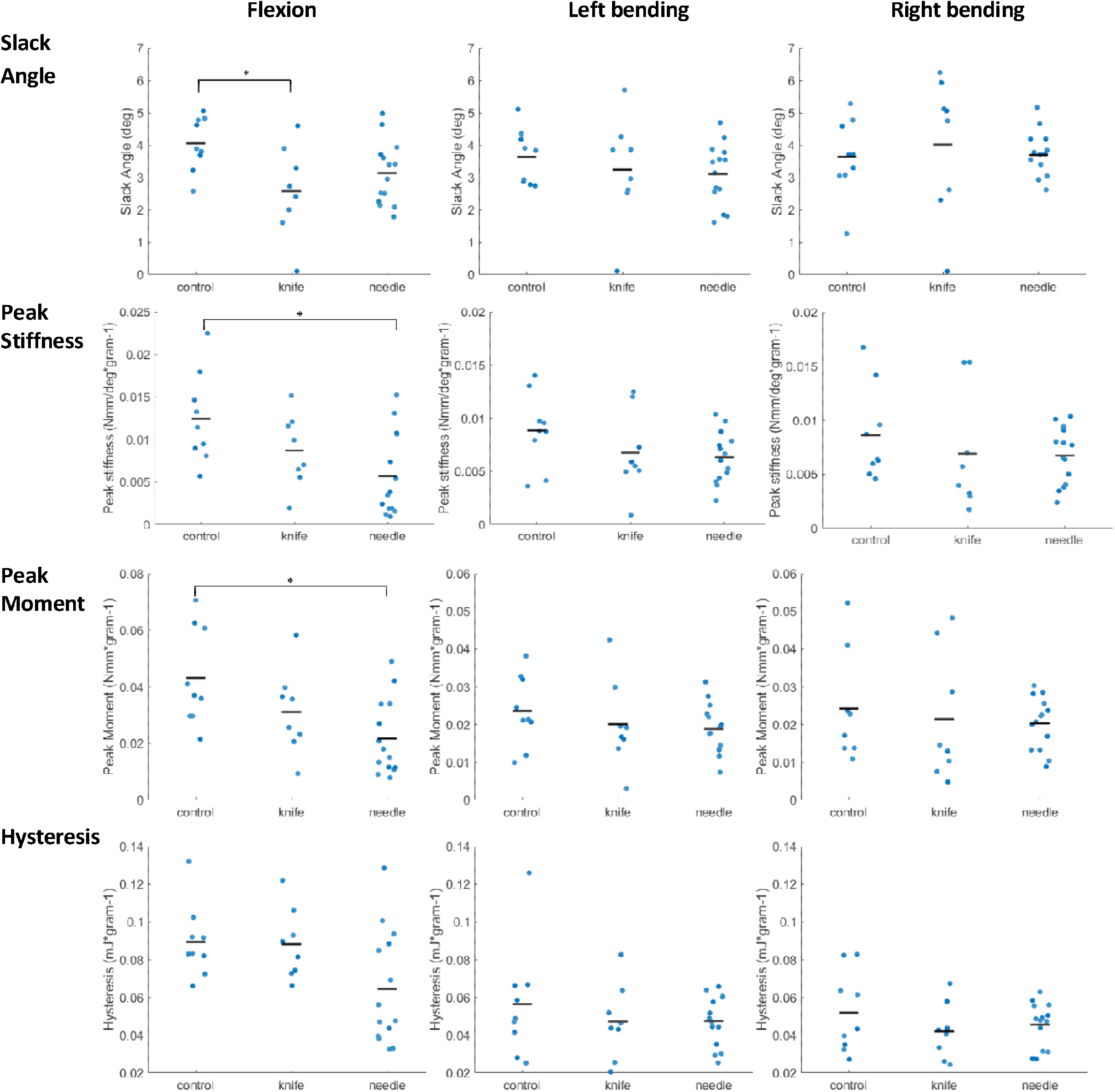
IVD mechanical properties. Dots represent individual data of each specimen, black bars represent the mean of the group. Asterisks (*) indicate a significant difference between groups.

**Table 3.** Mechanical properties of IVD.

| Group | sample size | Injury at L4/L5 | Slack Angle (deg) |  |  | Peak Stiffness (Nmm/deg*gram<sup>-1</sup>) |  |  | Peak Moment (Nmm*gram<sup>-1</sup>) |  |  | Hysteresis (mJ*gram<sup>-1</sup>) |  |  |
| --- | --- | --- | --- | --- | --- | --- | --- | --- | --- | --- | --- | --- | --- | --- |
|  |  |  | Flexion | Left bending | Right bending<sup>#</sup> | Flexion | Left bending | Right bending<sup>#</sup> | Flexion<sup>#</sup> | Left bending | Right bending<sup>#</sup> | Flexion | Left bending<sup>#</sup> | Right bending |
| control | 9 | NA | 4.06<br>(0.83) | 3.64<br>(0.85) | 3.72<br>(1.52) | 0.0125<br>(0.0053) | 0.0089<br>(0.0035) | 0.0064<br>(0.0036) | 0.0370<br>(0.0311) | 0.0236<br>(0.0094) | 0.0228<br>(0.0098) | 0.0895<br>(0.0192) | 0.0488<br>(0.0248) | 0.052<br>(0.0213) |
| knife | 8 | 8 | 2.58<br>(1.40) | 3.24<br>(1.63) | 4.91<br>(2.78) | 0.0087<br>(0.0042) | 0.0068<br>(0.0038) | 0.00485<br>(0.0059) | 0.0307<br>(0.0147) | 0.0201<br>(0.0117) | 0.0138<br>(0.0230) | 0.0883<br>(0.0187) | 0.0452<br>(0.0162) | 0.0421<br>(0.0149) |
| needle | 14 | 8<br>(2, L3/L4;<br>4, L5/L6) | 3.14<br>(0.98) | 3.11<br>(0.95) | 3.73<br>(0.97) | 0.0057<br>(0.0049) | 0.0063<br>(0.0024) | 0.00717<br>(0.0045) | 0.0165<br>(0.0207) | 0.0189<br>(0.0066) | 0.0215<br>(0.0109) | 0.0646<br>(0.0300) | 0.0481<br>(0.0221) | 0.0456<br>(0.0117) |

### IVD injury induced IVD structural changes

No group effects on IVD height and area were observed, but significant differences were found in absolute and relative NP area. The needle group exhibited smaller absolute and relative NP areas than the control group (*p*=0.020, 0.011, respectively), while no other post-hoc differences were found (Fig.3 & Table 1, 4, 5).

**Fig. 3.**
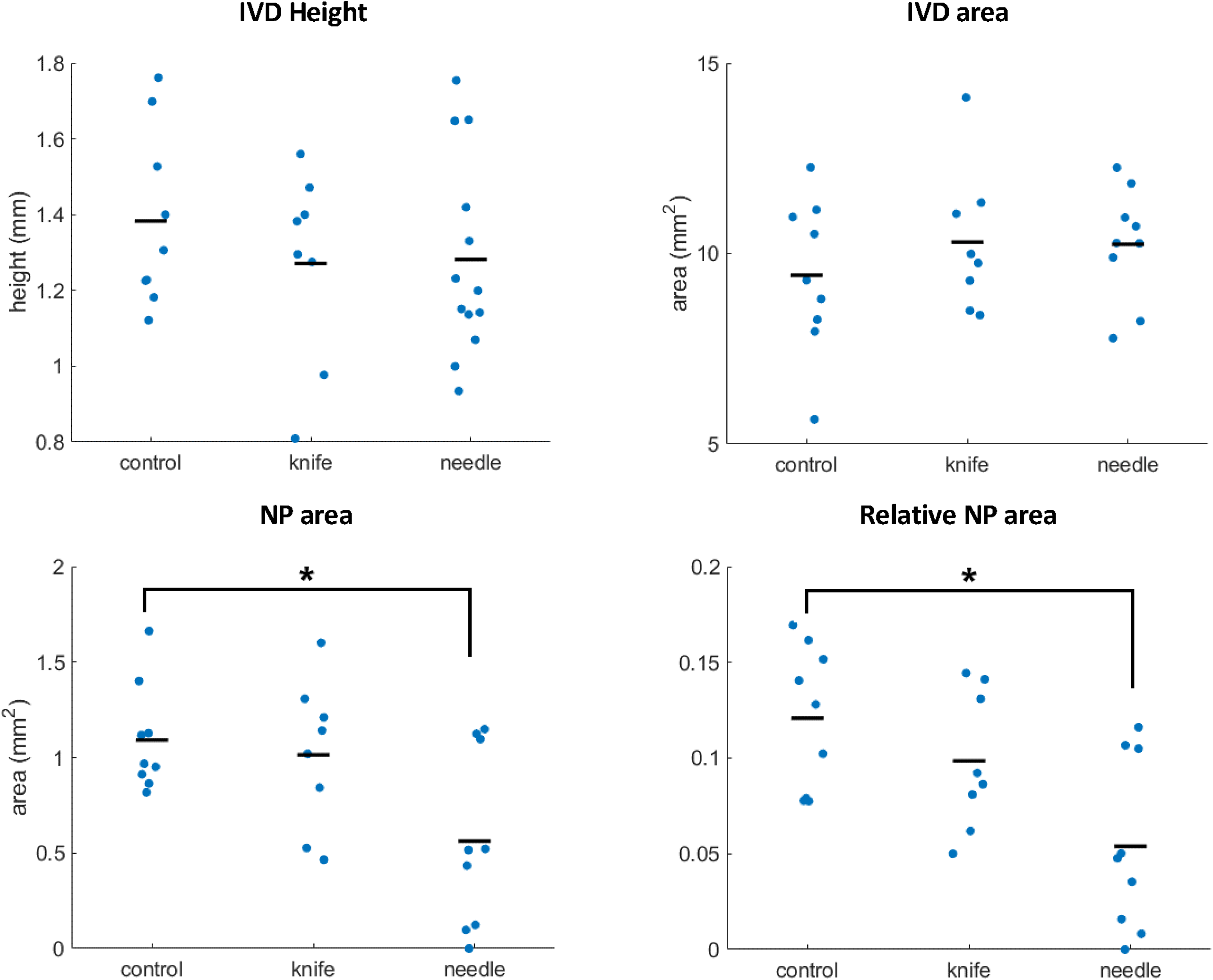
IVD structural parameters. Dots represent individual data of each specimen, black bars represent the mean of the group. Asterisks (*) indicate a significant difference between groups.

**Table 4.** Overview of ANOVA statistical results of IVD histology and mechanics.

| Variable | F (df) | P-value | Effect Size ($\eta^2$) | Pairwise Comparisons | t-value | p-value (Tukey adjusted) | Effect Size (Cohen's d) |
| --- | --- | --- | --- | --- | --- | --- | --- |
| IVD Height | 0.557 (2, 27) | 0.579 | 0.04 |  |  |  |  |
| IVD area | 0.643 (2, 23) | 0.535 | 0.053 |  |  |  |  |
| NP area | 4.969 (2, 23) | **0.016** | 0.302 | control – knife | 0.415 | 0.910 | 0.20 |
|  |  |  |  | control – needle | 2.934 | **0.020** | 1.38 |
|  |  |  |  | knife - needle | 2.432 | 0.058 | 1.18 |
| Slack Angle – Flexion | 4.218 (2, 28) | **0.025** | 0.232 | control – knife | 2.850 | **0.022** | 1.38 |
|  |  |  |  | control – needle | 2.010 | 0.128 | 0.86 |
|  |  |  |  | knife - needle | -1.180 | 0.473 | -0.52 |
| Slack Angle – Left bending | 0.614 (2, 28) | 0.548 | 0.042 |  |  |  |  |
| Peak Stiffness – Flexion | 5.348 (2, 28) | **0.011** | 0.276 | control – knife | 1.590 | 0.268 | 0.77 |
|  |  |  |  | control – needle | 3.260 | **0.008** | 1.40 |
|  |  |  |  | knife - needle | 1.410 | 0.350 | 0.62 |
| Peak Stiffness – Left bending | 1.866 (2, 28) | 0.173 | 0.118 |  |  |  |  |
| Peak Moment – Left bending | 0.799 (2, 28) | 0.460 | 0.054 |  |  |  |  |
| Hysteresis – Flexion | 3.735 (2, 28) | **0.036** | 0.211 | control – knife | 0.103 | 0.994 | 0.05 |
|  |  |  |  | control – needle | 2.362 | 0.064 | 1.01 |
|  |  |  |  | knife - needle | 2.164 | 0.095 | 0.96 |
| Hysteresis – Right bending | 0.888 (2, 28) | 0.423 | 0.06 |  |  |  |  |
| Normalized MF mass | 1.166 (2, 27) | 0.327 | 0.079 |  |  |  |  |
| Normalized ML mass | 5.24 (2, 26) | **0.012** | 0.287 | control – knife | -0.234 | 0.970 | -0.12 |
|  |  |  |  | control – needle | -2.731 | **0.029** | -1.26 |
|  |  |  |  | knife - needle | -2.580 | **0.041** | -1.14 |

*IVD, intervertebral disc; NP, nucleus pulposus; MF, multifidus muscle; ML, longissimus muscle; df, degree of* *freedom*

**Table 5.**
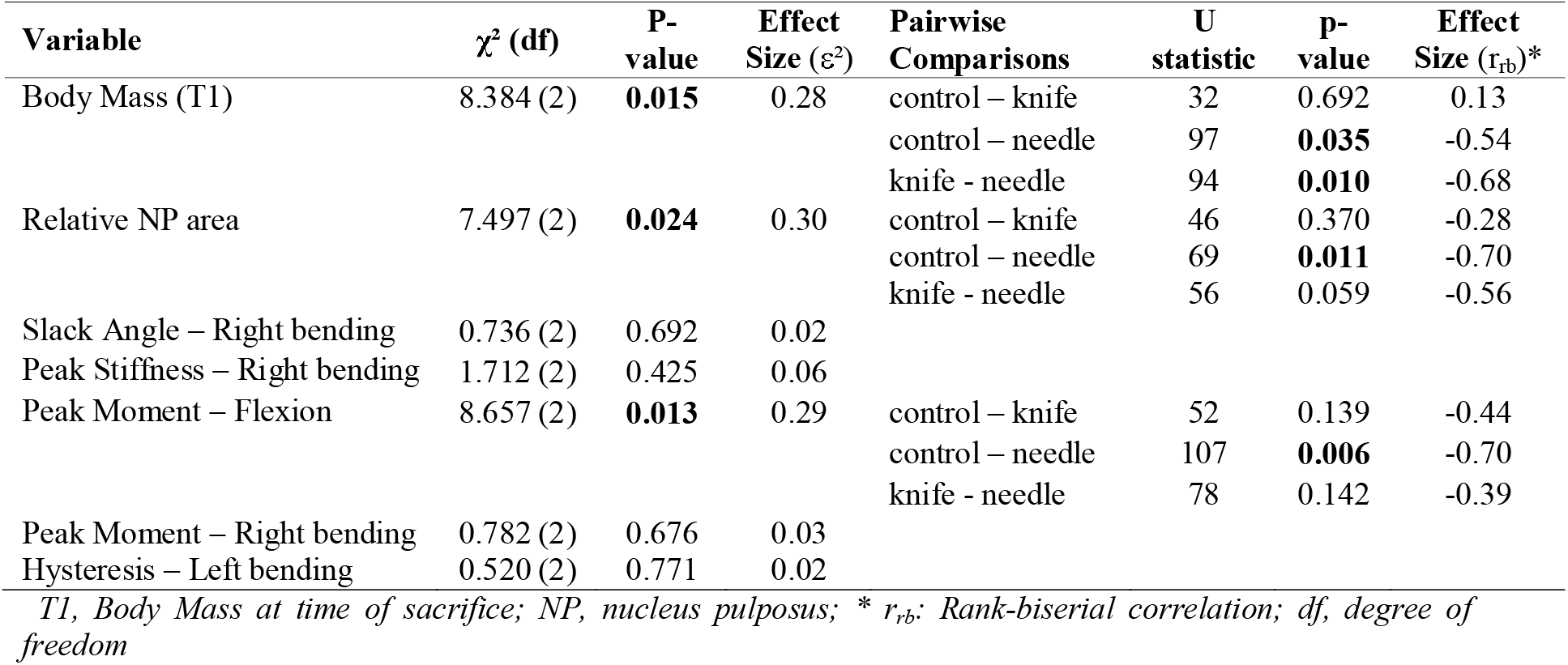
Overview of Kruskal-Wallis statistical results of IVD histology and mechanics.

PR staining (Fig.4) showed intact and well-organized AF fiber rings, a round-shaped NP, and a clear border between AF and NP in the control group. In the knife group, the AF fiber rings were less organized, NP was less round, and the border was less clear. In the needle group, the AF rings were also less organized and even ruptured, the NP lacked a round shape, the border between AF and NP was unclear, and the needle tract was visible in some specimens.

**Fig. 4.**
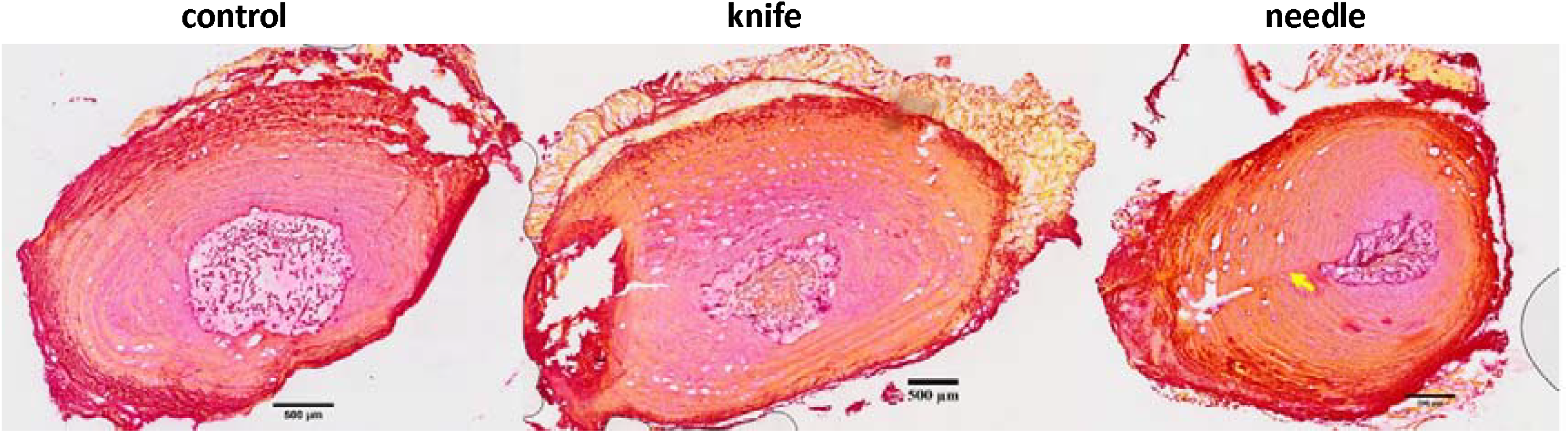
Examples of sections of the intervertebral disc stained with Picrosirius Red. Less organized annuls fibrosus (AF), less round-shaped nucleus pulposus (NP), and less clear border between AF and NP were observed in the knife and needle groups. Yellow arrow points to the needle puncture tract. Scale bar=500 µm

### Muscle properties

No group effect was found in normalized MF mass, but a significant group effect was found in normalized ML mass (*p*=0.012, F_(2,26)_=5.24), with the needle group having a significantly higher mass than the control (*p*=0.029, t=-2.731) and knife groups (*p*=0.041, t=-2.580) (Fig.5 & Table 1, 4).

**Fig. 5.**
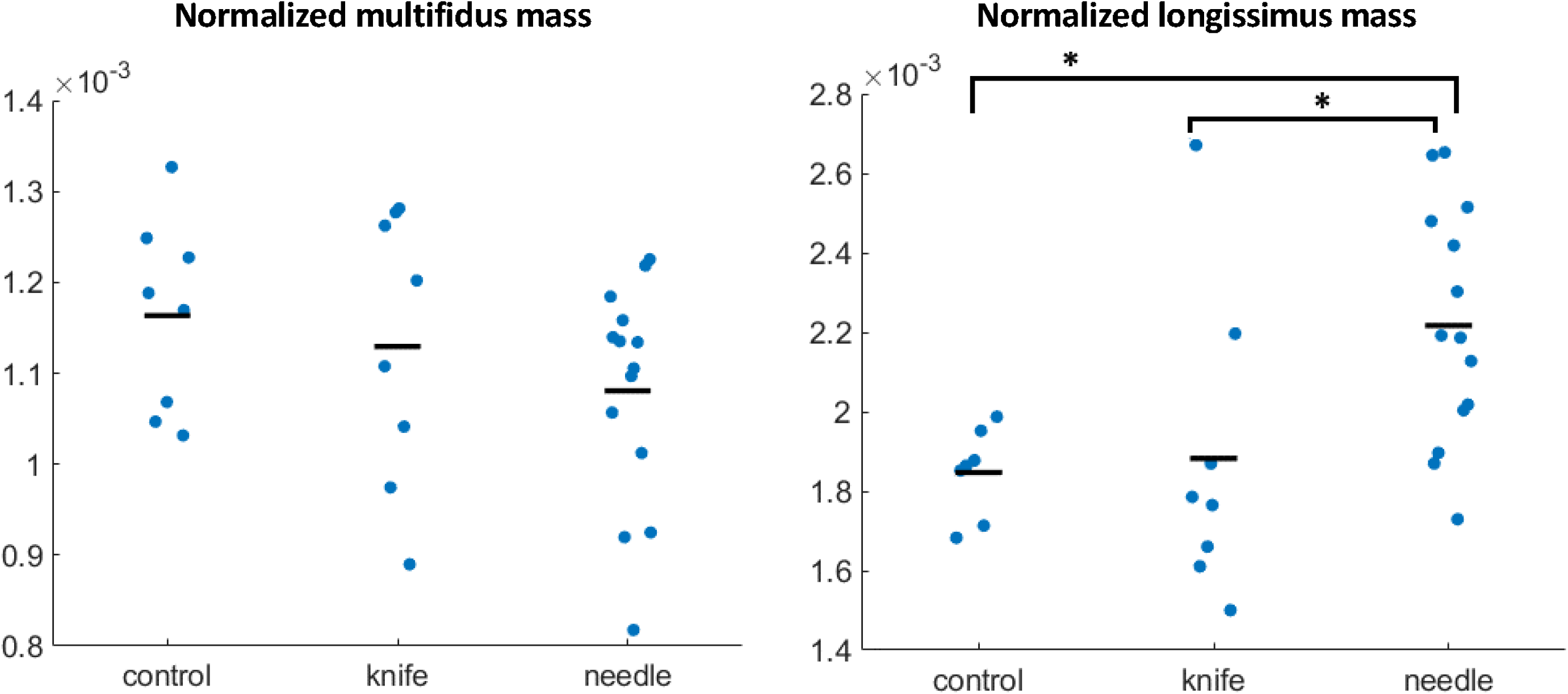
Normalized mass of multifidus (left) and longissimus (right) muscle. Dots represent individual data of each specimen, black bars represent the mean of the group. Asterisks (*) indicate a significant difference between groups.

## Discussion

Distinct mechanical and structural consequences were observed within one week after IVD injury, in particular after needle injury, supported that this intervention provides a successful rat model of lumbar spine instability. IVD injury, using a 21G needle, caused significant mechanical instability in flexion and morphological degeneration, as evidenced by decreased IVD mechanical properties such as peak stiffness, peak moment, and hysteresis, and reduced NP size, morphological degeneration. However, IVD height and area remained unchanged. No significant effects of IVD injury on left and right bending mechanics were observed. While structural degeneration was evident through histological analysis. With respect to muscle adaptation, there was no significant alteration in normalized MF mass, but an increased normalized ML mass was found in the needle group.

Various methods have been used to induce IVD degeneration in rodent models, with degeneration severity affected by parameters, such as injury depth (Han et al. 2008), number (Ulrich et al. 2007) and size (Elliott et al. 2008; Martin et al. 2013). In this study, we compared knife stab lesion and needle puncture. Our results indicate that needle puncture changed mechanical properties of the IVD, specifically, peak stiffness (54%) and peak moment (55%) in flexion, comparable to the 60% reduction of compressive and torsional stiffness reported after mouse tail disc injuries using a 26G needle (Martin et al. 2013). Despite the fact that the knife’s maximum thickness (equal to 43% of the average disc height in the control group) exceeded the suggested 40% threshold for inducing mechanical changes (Elliott et al. 2008), no significant effects on IVD mechanics were observed. Possibly the needle puncture created more damage to the IVD compared to the knife stab lesion. Firstly, the thickness of the knife decreases towards the tip, so even with a relatively large width, the damage in the center of the IVD might not be sufficient. In addition, the needle removed some disc tissue when rotated 360° before withdrawal. More NP tissue may have herniated through the needle tract, causing more depressurization and shrinkage of the NP (Fig.3), which could be confirmed by the smaller NP size in the needle group than in the knife group (Table 1). These differences may have been increased by partial healing of the AF through granulation tissue formation (Kaapa et al. 1995) or spontaneous repair (Schollum et al. 2010). In smaller defects, this repair process might close the annular tear, allowing NP re-pressurization via nuclear matrix synthesis, thereby restoring disc mechanical properties. In contrast, larger defects may require longer wound closure time and prolong the impaired disc mechanical function (Lotz 2004).

Our study quantitatively and qualitatively assessed the effects of injury on IVD morphology. In contrast to previous studies in rats, no significant injury effects on IVD height were found, with only a 7-8% height loss after injury, compared to approximately 20% height reduction after lumbar (Mosley et al. 2020) and tail (Zhu et al. 2023) IVD injuries. Compared to the control group, we observed a similar IVD area, but a smaller NP area in the injury groups, consistent with findings from rat (Zhang et al. 2009; Maas et al. 2018), porcine (Niinimaki et al. 2007), and rabbit (Jacobs et al. 2013) IVD injury studies. Specifically, the relative NP area in the knife and needle groups showed 30% and 63% reductions, respectively, consistent with our previous findings (Maas et al. 2018). Histological staining confirmed degenerative changes similar to those in previous studies, including disorganized AF fiber rings, fiber rupture, decreased NP area in the injured discs, as well as loss of a distinct border between AF and NP (Mosley et al. 2020; Zhu et al. 2023). These findings indicate that our injury was sufficient to induce morphological degeneration of the IVD.

With respect to muscular responses to IVD injury, no significant effects on normalized MF mass were observed in our study. Our previous study also found no change in MF fiber type composition following the same lumbar IVD knife stab lesion in the rat (Docter et al. 2021). However, MF atrophy within one week following lumbar IVD injury, indicated by decreased normalized mass (Maas et al. 2018) and cross-sectional area (Hodges et al. 2006) has been reported. Interestingly, we found an increase in normalized ML mass, suggesting a compensatory adaptation, to maintain spinal stability. Further research is needed to explore the specific responses of the paraspinal muscles, for instance, changes in connective tissue content and mechanical properties.

One limitation of this paper is that the control group animals were one week older than those in the injury groups. However, previous studies observed no significant changes in IVD structure (Maas et al. 2018), mechanics (Martin et al. 2013), or back muscle mass (Maas et al. 2018) within one week. Our method for assessing disc height, although practical for this study, is not as accurate as MRI (Li et al. 2019) or X-ray (Mosley et al. 2020). For the needle group, the level of injury was not L4/L5 only. As IVD area (Maas et al. 2018) and height are similar for the adjacent levels (confirmed by similar means in Table 1), this is not expected to cause bias. In addition, this study only assessed changes within one week post-injury. Degenerative changes in the IVD and muscular adaptations may take longer to fully manifest (Maas et al. 2018). A longer follow-up period would provide more insight into the chronic effects of IVD injury.

Our findings emphasize the complexity of the IVD’s response to injury. However, the increase in normalized ML mass suggests that muscular adaptations may occur as a compensatory response soon after IVD lesion, providing insights into the neuromuscular consequences of IVD degeneration. This study holds potential clinical relevance by improving our understanding of how IVD injury and degeneration contribute to spinal instability.

## Acknowledgements

We would like to thank Guus Baan for making the illustrations for surgery (Fig.1A) and mechanical test setup (Fig.1B).

