## supplemental Table 1-2 for "An in vivo rat lumbar spine instability model induced by intervertebral disc injury"

**Appendix A**

| **Table 1. Normality Test (Shapiro-Wilk)** | | |
| --- | --- | --- |
|  | **W** | **p** |
| Body Mass (T1) | 0.903 | 0.009 |
| IVD Height | 0.972 | 0.585 |
| IVD area | 0.995 | 1.000 |
| NP area | 0.935 | 0.100 |
| Relative NP area | 0.909 | 0.025 |
| Slack Angle – Flexion | 0.987 | 0.959 |
| Slack angle – Left bending | 0.968 | 0.476 |
| Slack angle – Right bending | 0.952 | 0.181 |
| Peak Stiffness – Flexion | 0.935 | 0.059 |
| Peak Stiffness – Left bending | 0.971 | 0.549 |
| Peak Stiffness – Right bending | 0.904 | 0.009 |
| Peak Moment – Flexion | 0.922 | 0.027 |
| Peak Moment – Left bending | 0.977 | 0.736 |
| Peak Moment – Right bending | 0.917 | 0.020 |
| Hysteresis – Flexion | 0.932 | 0.051 |
| Hysteresis – Left bending | 0.913 | 0.015 |
| Hysteresis – Right bending | 0.954 | 0.197 |
| Normalized mass MF | 0.937 | 0.076 |
| Normalized mass LG | 0.964 | 0.421 |
| *T1, Body Mass at time of termination; IVD, intervertebral disc; NP, nucleus pulposus; MF, multifidus muscle; LG, longissimus muscle* | | |

| **Table 2. Homogeneity of Variances Test (Levene's)** | | | | |
| --- | --- | --- | --- | --- |
|  | **F** | **df1** | **df2** | **p** |
| Body Mass (T1) | 2.218 | 2 | 28 | 0.128 |
| IVD Height | 0.121 | 2 | 27 | 0.886 |
| IVD area | 0.547 | 2 | 23 | 0.586 |
| NP area | 1.433 | 2 | 23 | 0.259 |
| Relative NP area | 0.305 | 2 | 23 | 0.740 |
| Slack Angle – Flexion | 0.853 | 2 | 28 | 0.437 |
| Slack angle – Left bending | 1.34 | 2 | 28 | 0.278 |
| Slack angle – Right bending | 7.61 | 2 | 28 | 0.002 |
| Peak Stiffness – Flexion | 0.195 | 2 | 28 | 0.824 |
| Peak Stiffness – Left bending | 0.606 | 2 | 28 | 0.552 |
| Peak Stiffness – Right bending | 2.283 | 2 | 28 | 0.121 |
| Peak Moment – Flexion | 0.544 | 2 | 28 | 0.587 |
| Peak Moment – Left bending | 0.819 | 2 | 28 | 0.451 |
| Peak Moment – Right bending | 4.594 | 2 | 28 | 0.019 |
| Hysteresis – Flexion | 3.074 | 2 | 28 | 0.062 |
| Hysteresis – Left bending | 1.301 | 2 | 28 | 0.288 |
| Hysteresis – Right bending | 3.688 | 2 | 28 | 0.038 |
| Normalized mass MF | 0.872 | 2 | 27 | 0.430 |
| Normalized mass LG | 2.854 | 2 | 26 | 0.076 |
| *T1, Body Mass at time of termination; IVD, intervertebral disc; NP, nucleus pulposus; MF, multifidus muscle; LG, longissimus muscle; df, degree of freedom* | | | | |
