## supplemental Fig.1 for "An in vivo rat lumbar spine instability model induced by intervertebral disc injury"

**Appendix B**

| **A** | 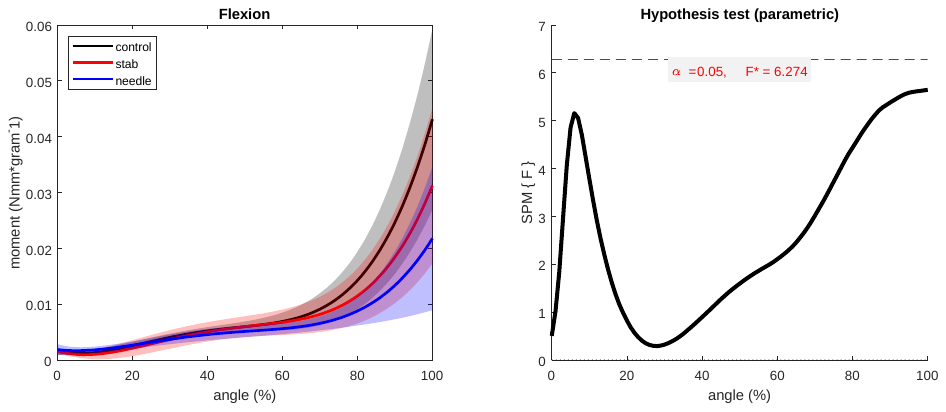 |
| --- | --- |
| **B**.1 | 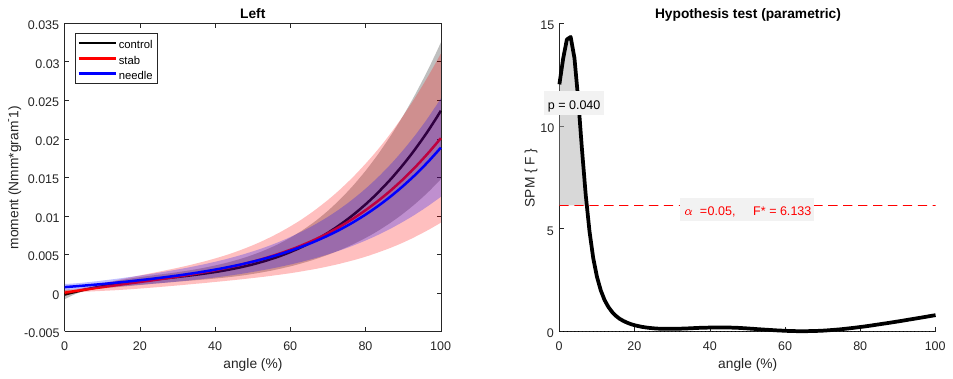 |
| **B**.2 | 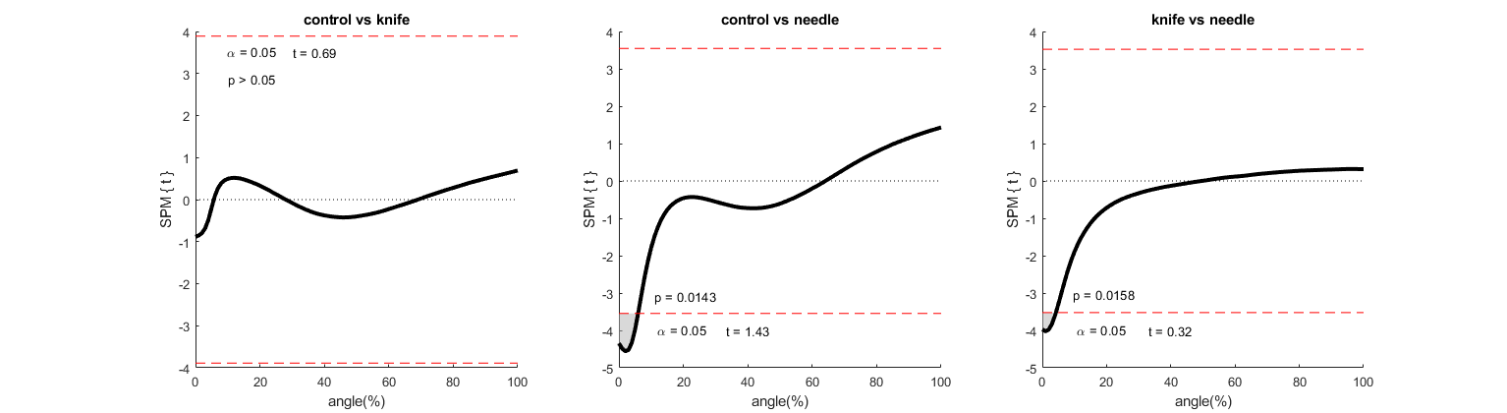 |
| **C.**1 | 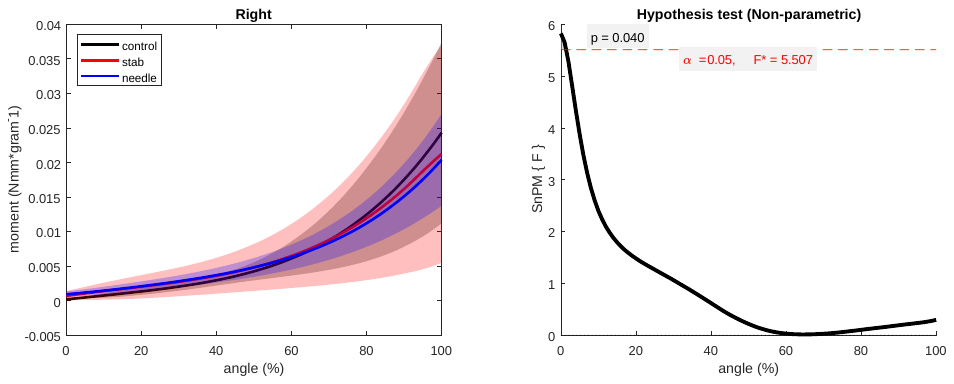 |
| **C.**2 | 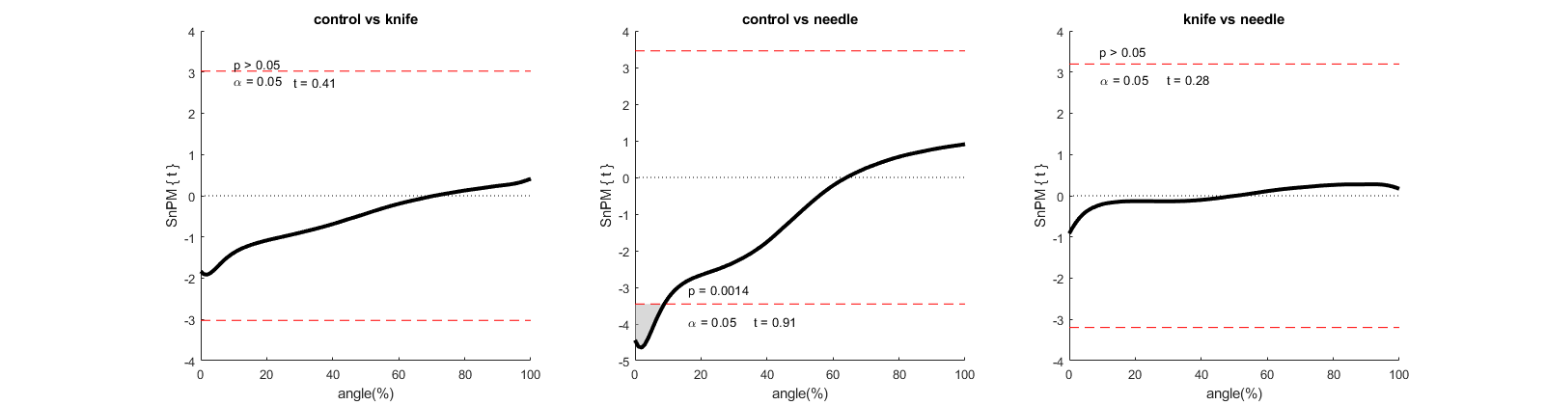 |
| **Fig.1**. Comparison of angle-moment curves in **(A)** flexion, **(B**.1**)** left bending and **(C.**1**)** right bending. Moments are plotted as a function of normalized bending angle and presented as mean with 95 % confidence interval (shaded area). Pairwise comparison of different groups in left bending (**B**.2) and right bending (**C**.2). The critical threshold was set as α=0.05 (red dashed line). Gary zone indicated region with statistically significant difference. | |
